# A computational framework for systematic exploration of biosynthetic diversity from large-scale genomic data

**DOI:** 10.1101/445270

**Authors:** Jorge C. Navarro-Muñoz, Nelly Selem-Mojica, Michael W. Mullowney, Satria Kautsar, James H. Tryon, Elizabeth I. Parkinson, Emmanuel L.C. De Los Santos, Marley Yeong, Pablo Cruz-Morales, Sahar Abubucker, Arne Roeters, Wouter Lokhorst, Antonio Fernandez-Guerra, Luciana Teresa Dias Cappelini, Regan J. Thomson, William W. Metcalf, Neil L. Kelleher, Francisco Barona-Gomez, Marnix H. Medema

## Abstract

Genome mining has become a key technology to explore and exploit natural product diversity through the identification and analysis of biosynthetic gene clusters (BGCs). Initially, this was performed on a single-genome basis; currently, the process is being scaled up to large-scale mining of pan-genomes of entire genera, complete strain collections and metagenomic datasets from which thousands of bacterial genomes can be extracted at once. However, no bioinformatic framework is currently available for the effective analysis of datasets of this size and complexity. Here, we provide a streamlined computational workflow, tightly integrated with antiSMASH and MIBiG, that consists of two new software tools, BiG-SCAPE and CORASON. BiG-SCAPE facilitates rapid calculation and interactive visual exploration of BGC sequence similarity networks, grouping gene clusters at multiple hierarchical levels, and includes a ‘glocal’ alignment mode that accurately groups both complete and fragmented BGCs. CORASON employs a phylogenomic approach to elucidate the detailed evolutionary relationships between gene clusters by computing high-resolution multi-locus phylogenies of all BGCs within and across gene cluster families (GCFs), and allows researchers to comprehensively identify all genomic contexts in which particular biosynthetic gene cassettes are found. We validate BiG-SCAPE by correlating its GCF output to metabolomic data across 403 actinobacterial strains. Furthermore, we demonstrate the discovery potential of the platform by using CORASON to comprehensively map the phylogenetic diversity of the large detoxin/rimosamide gene cluster clan, prioritizing three new detoxin families for subsequent characterization of six new analogs using isotopic labeling and analysis of tandem mass spectrometric data.

## Introduction

Microbial specialized metabolites are key mediators of interspecies communication and competition in the environment and in the context of host microbiomes^1,2^. Their diverse chemical structures have been critical in the development of antibiotics, anticancer drugs, crop protection agents and ingredients for manufacturing. While tens of thousands of natural products have been discovered in past decades, recent evidence suggests that these represent a fraction of the potential natural product chemical space yet to be discovered^3–8^.

Genome mining has emerged in the past decade as a key technology to explore and exploit natural product diversity. Key to this success is the fact that genes encoding natural product biosynthetic pathways are usually clustered together on the chromosome. These biosynthetic gene clusters (BGCs) can be readily identified in a genome. Moreover, in many cases, the chemical structures of their products can be predicted to a certain extent, based on the analysis and biosynthetic logic of the enzymes encoded in a BGC and their similarity to known counterparts^9^.

Initially, genome mining was performed on a single-genome basis: a research group or consortium would sequence the genome of a single microbial strain and attempt to identify and characterize each of its BGCs one by one. This approach has revealed much about the metabolic capacities of model natural-product-producing organisms like *Streptomyces coelicolor*, *Sorangium cellulosum* and *Aspergillus nidulans* and has provided clues regarding the discovery potential from corresponding genera^10–12^. Computational tools for the identification of BGCs and the prediction of their products’ chemical structures, such as antiSMASH^13–16^ and PRISM^17–19^, have played a key role in the success of genome mining. These *in silico* approaches have been strengthened by comparative analysis of identified BGCs with rich empirical reference data, such as those provided by the MIBiG community effort, which has documented (meta)data on >1,000 BGCs connected to their products^20^.

Fueled by rapid developments in high-throughput sequencing, genome mining efforts are now expanding to large-scale pan-genomic mining of entire bacterial genera^4,21,22^, strain collections^23^ and metagenomic datasets from which thousands of metagenome-assembled genomes (MAGs) can be extracted at once^24–27^. Such studies pave the path towards systematic investigations of the biosynthetic potential of broad taxonomic groups of organisms, as well as entire ecosystems. These large-scale analyses easily lead to the identification of thousands of BGCs with varying degrees of mutual similarity, ranging from widely distributed homologs of gene clusters for the production of well-known molecules to rare or unique gene clusters that encode unknown enzymes and pathways.

To map and prioritize this complex biosynthetic diversity, several groups have devised methods to compare architectural relationships between BGCs in sequence similarity networks and group them into gene cluster families (GCFs), each of which contains BGCs across a range of organisms that are linked to a highly similar natural product chemotype^4,28,29,^. The presence or expression of such GCFs can be correlated to molecular families (MFs) identified from mass spectrometry data to match genes to their product molecules in a process termed metabologenomics^4,30,31^. However, current methods fail to correctly measure similarity between complete and fragmented gene clusters (which frequently occur in metagenomes and large-scale pan-genome sequencing projects based on short-read technologies); do not consider the complex and multi-layered evolutionary relationships within and between GCFs; require lengthy CPU-time on supercomputers when processing large datasets; and lack a user-friendly implementation that interacts directly with other key resources. These shortcomings preclude adoption by the broader scientific community and impede significant advances in natural product discovery.

Here, we provide a streamlined computational workflow that tightly integrates two new software tools, BiG-SCAPE and CORASON, with the gene cluster identification and empirical biosynthetic data comparison possible through antiSMASH^16^ and MIBiG^20^ (Fig 1). BiG-SCAPE facilitates rapid calculation and interactive exploration of BGC sequence similarity networks (SSNs); it accounts for differences in modes of evolution between BGC classes, groups gene clusters at multiple hierarchical levels (families, clans and classes), introduces a ‘glocal’ alignment mode that supports complete as well as fragmented BGCs, and democratizes the analysis through a dramatically accelerated and interactive stand-alone user interface. As a complement to this, CORASON employs a phylogenomic approach to elucidate detailed evolutionary relationships between gene clusters by computing high-resolution multi-locus phylogenies of all BGCs within and across GCFs. Additionally, it allows researchers to comprehensively identify all genomic contexts in which gene cassettes of interest, which encode a specific function within a BGC, can be found. To validate the GCF classifications, we show that metabologenomic correlations accurately connect genes to mass features across metabolomic data from 363 strains. Furthermore, we demonstrate the power of the combined workflow, together with the EvoMining algorithm^8^, to comprehensively map diversity within gene cluster clans by identifying three new families responsible for the biosynthesis of new detoxins.

**Fig 1.**
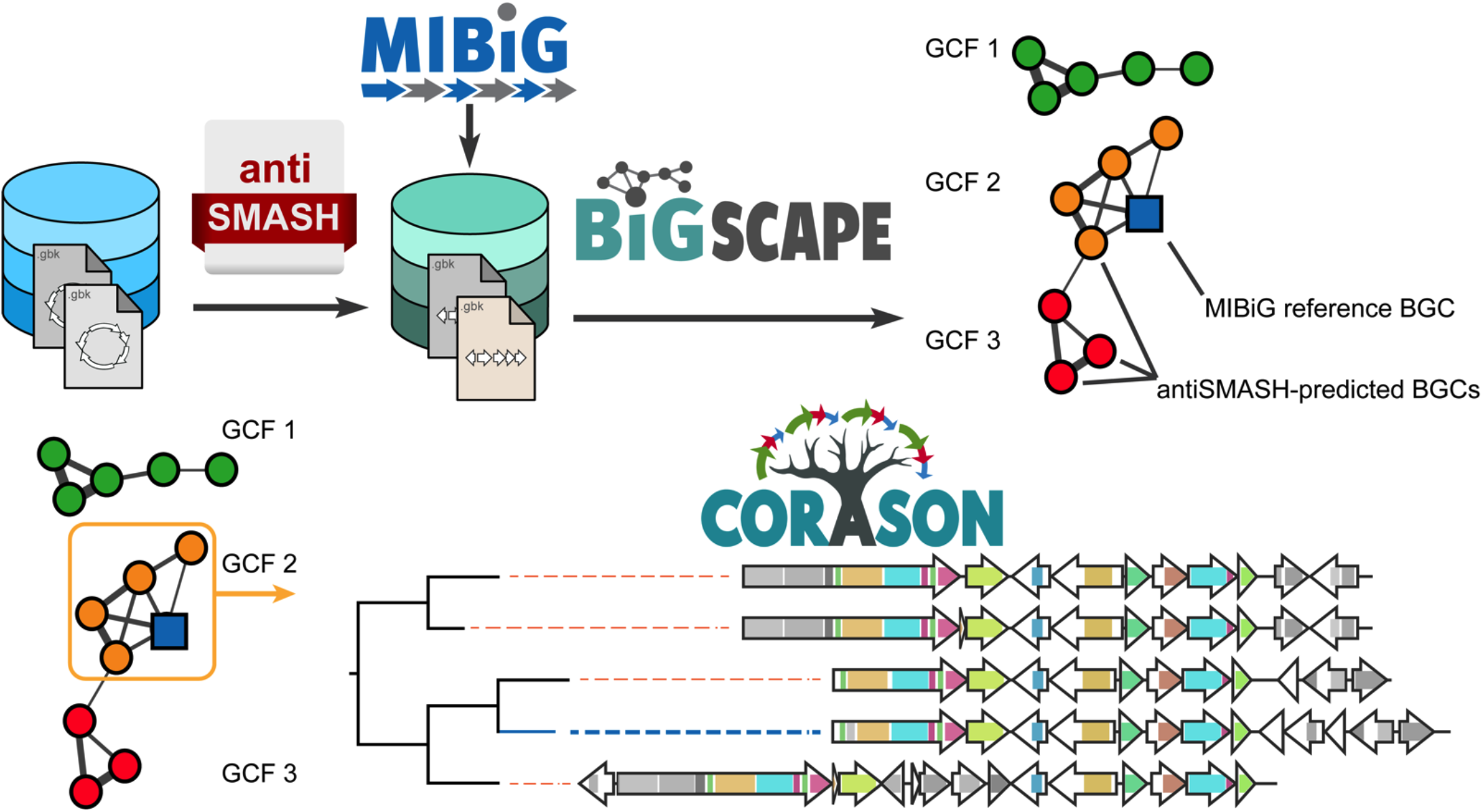
The BiG-SCAPE/CORASON workflow. The BiG-SCAPE approach analyzes a set of biosynthetic gene clusters to construct a BGC similarity network and groups them into GCFs. Subsequently, CORASON-based multi-locus phylogenetic analysis is employed to illuminate evolutionary relationships of BGCs within each GCF.

## Results & Discussion

### A streamlined algorithm for large-scale network analysis and classification of biosynthetic gene clusters

To provide a streamlined, scalable and user-friendly software for exploring and classifying large collections of gene clusters, we built the Biosynthetic Gene Similarity Clustering and Prospecting Engine (BiG-SCAPE), written in Python and freely available as open source software (https://git.wageningenur.nl/medema-group/BiG-SCAPE). BiG-SCAPE takes BGCs predicted by antiSMASH or annotated in MIBiG as inputs to automatically generate sequence similarity networks and assemble GCFs.

In previous studies by Cimermancic et al.^3^ and Doroghazi et al.^4^, two sets of distance metrics had been independently developed to measure the (dis)similarity of pairs of BGCs. In BiG-SCAPE, we aimed to combine the respective strengths of both approaches. The strength of the former approach was the elegant compression of gene clusters into strings of Pfam domains, combined with the Jaccard index (JI) to measure domain content similarity (Fig 2a). However, in the metric of Cimermancic et al.^3^, a useful index for synteny conservation had been missing; to this end, we added an Adjacency Index (AI), which measures how many pairs of adjacent domains are shared between gene clusters.

**Fig. 2.**
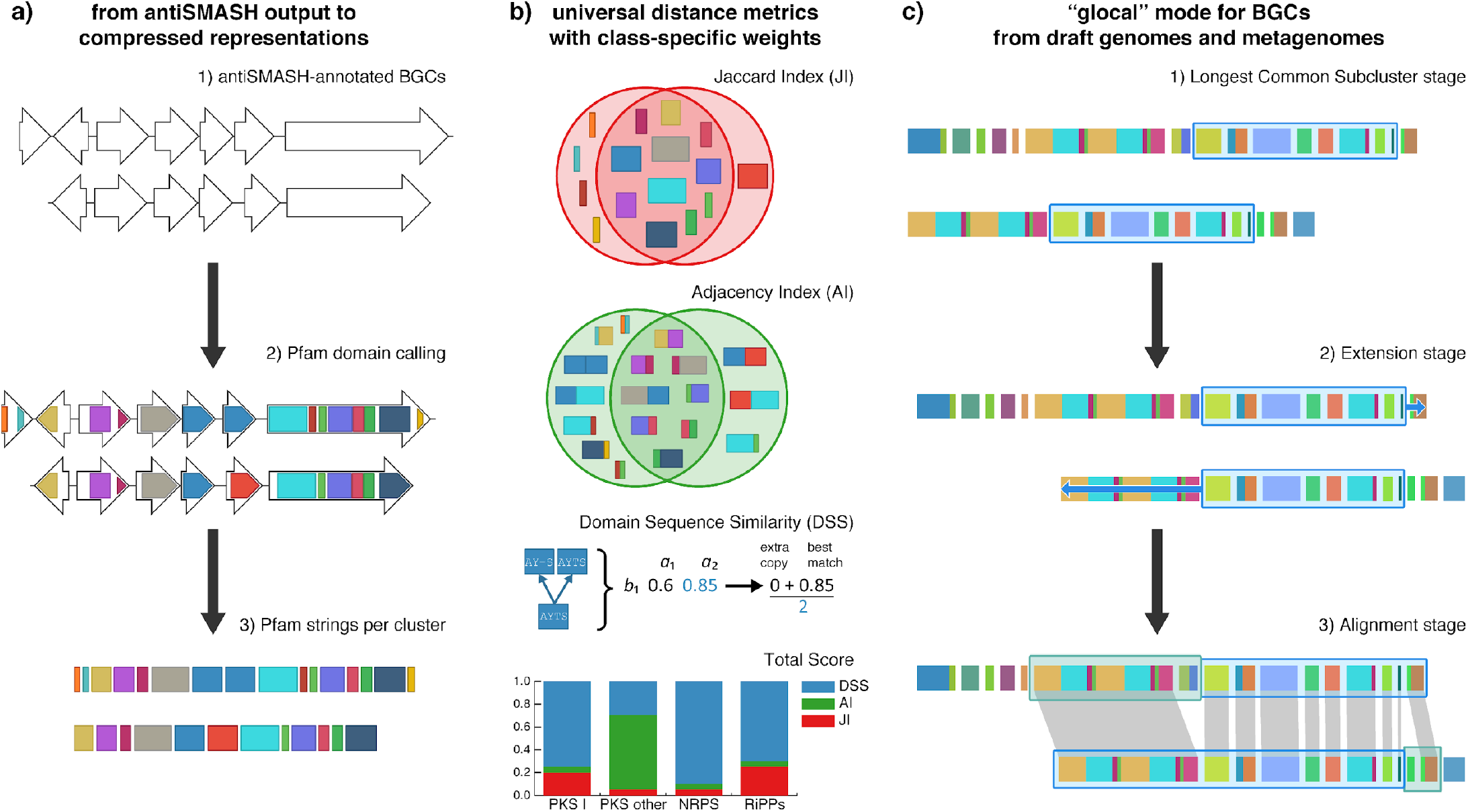
Main concepts in the BiG-SCAPE algorithm. **a,** Input data comes from the genomic loci of BGCs, which are directly imported from antiSMASH and/or MIBiG. Nucleotide sequences are translated and represented as strings of Pfam domains; **b,** The three metrics that are combined in a single distance include the Jaccard Index (JI), which measures the percentage of shared types of domains; the Adjacency Index (AI), which measures the percentage of pairs of adjacent domains; and the Domain Sequence Similarity (DSS), which measures the identity of profile-aligned sequences in the conserved domains (details in Text S1); **c,** in “glocal” mode, BiG-SCAPE starts with the longest common subcluster of genes between a pair of BGCs and attempts to extend the selection of genes for comparison.

Finally, sequence identity is an important parameter, as Pfam domains are often very broad and frequently comprise a wide range of enzyme subfamilies with different catalytic activities or substrate specificities. Yet, all previously developed approaches suffered from extremely long compute times when including sequence identity calculations, requiring the use of supercomputers that would preclude day-to-day use by natural product scientists worldwide. The underlying issue is that comparing large numbers of protein sequences from many BGCs is an all-versus-all problem that scales quadratically when the size of the data increases. To mitigate this, we replaced all-versus-all calculations with all-versus-profile calculations, by aligning each protein domain sequence to its profile Hidden Markov Model from Pfam using the hmmalign tool^32^. This leads to a marked speed increase compared to conventional multiple sequence alignment using Muscle: when calculating a network of the 1,393 BGCs in MIBiG version 1.3, an implementation with hmmalign took 15 minutes, while a Muscle implementation took 223 minutes [see online methods: Alignment Method Comparison]. With larger numbers of BGCs, the differences are considerably larger. We implemented the profile-based alignment into the domain sequence similarity (DSS) index, an updated version of the domain duplication index, which measures both Pfam domain copy number differences and sequence identity. The combination of JI, AI and DSS indices into a new combined metric constitutes a fast and informative method to calculate distances between BGCs.

One notable limitation of a generic distance metric is that different classes of BGCs have different evolutionary dynamics. For example, the chemical structures of aryl polyenes have been shown to remain very stable across large evolutionary timescales, while the amino acid sequence identity between their key biosynthetic enzymes has become less than 30-40%^3^. On the other hand, the structures of rapamycin family polyketides exhibit major differences even when sequence identities are as high as ~80%^33^. Although there is not enough information available to construct individual metrics for each specific natural product family, we did calibrate specific weights of the JI, AI and DSS indices for BGCs encoding eight different BiG-SCAPE classes: type I polyketide synthases (PKS), other PKSs, nonribosomal peptide synthetases (NRPS), PKS/NRPS hybrids, RiPPs, saccharides, terpenes and others (see Fig. 2b and Table S5). In the output, separate networks are generated for each BiG-SCAPE class, along with an optional overall network that combines BGCs from all classes.

Another problem of previous approaches for calculating distances between BGCs was how to handle comparisons between complete and partial BGCs (e.g., from fragmented genome assemblies), as well as comparisons with pairs of genomically adjacent BGCs that are merged into one cluster by antiSMASH or other cluster identification tools. Both global similarity (used in all previous methods) and local similarity lead to artifacts in such cases. To compare the appropriate corresponding regions between BGCs, we introduced a new ‘glocal’ alignment mode, which first uses a fast algorithm to find the longest common substring between the Pfam strings of a BGC pair, and then uses match/mismatch penalties to extend the alignment until the end of the matching region (Fig. 2c, Online Methods). Information about whether an antiSMASH^16^-annotated cluster is located at the edge of a contig can also be used to automatically select a third pairwise distance calculation mode that relies on global alignment for complete clusters and glocal alignment when at least one of the BGCs in a pair is fragmented.

BGC sequence similarity networks are then generated by applying a cut-off to the distance matrix calculated by BiG-SCAPE, while rounds of affinity propagation clustering^34^ are used to group BGCs into GCFs, and GCFs into “Gene Cluster Clans” (GCCs). This process of categorization facilitates calculating metabologenomic correlations^35,36^ at multiple levels.

Results can be further processed in downstream analysis or immediately visualized using an interactive HTML-based interface that permits dynamic exploration of the gene cluster network (see further below for more details).

### Validation using natural product chemical similarity networks and large-scale metabolomics data

To verify that BiG-SCAPE is able to group together BGCs that are known to be related, we constructed a chemical similarity network from all products of BGCs in MIBIG, and used this to derive a curated set of 376 compounds, which were manually classified into 92 groups (e.g. 14-membered macrolides, benzoquinone ansamycins, quinomycin antibiotics etc.) and 9 classes (e.g. Polyketides, NRPs, RiPPs etc.). We then used BiG-SCAPE to group the corresponding BGCs into GCFs and observed good correspondence between manually curated families and those predicted by BiG-SCAPE (Supplementary Fig S5).

Arguably, the greatest value of BiG-SCAPE lies in the practical utility of the predicted GCFs for discovery applications. Hence, we assessed the accuracy of correlations of BiG-SCAPE-predicted GCFs to MS ions from known natural product through metabologenomics^31^. First, we performed a BiG-SCAPE analysis of 74,652 BGCs from 3,080 actinobacterial genomes (see Methods), including 1,393 reference BGCs from MIBiG^20^. BiG-SCAPE grouped these BGCs into a total number of 17,718 GCFs and 801 GCCs using default parameters. Extracts from 363 actinomycete strains were analyzed using untargeted high-resolution LC-MS/MS^4^. The GCF annotations for these 363 strains from two BiG-SCAPE modes (global and glocal) at two similarity cutoffs (0.30 and 0.50) were used to generate four rounds of metabologenomic correlations utilizing a binary scoring metric as described previously^4,30^. BiG-SCAPE’s gene cluster family annotations were then assessed against ion production patterns via two methods. First, a ‘golden dataset’ of nine known ion signals and their characterized gene clusters were manually tracked across the four correlation rounds. These ion signals corresponded to the following natural products; CE-108, benarthin, desertomycin, tambromycin, enterocin, tyrobetaine, chlortetracycline, rimosamide, and oxytetracycline. Second, a target-decoy approach was applied to estimate the false discovery rate (FDR) for each round to provide an overview of correlative power for the unknown ion to gene cluster family hypotheses generated (see Fig. S14). Decoy databases used for the target-decoy approach were made for each round by randomizing the Boolean arrays for both ion detection patterns and GCF member detection patterns. Correlations run with glocal mode GCF annotations outperformed global mode correlations, with 4/9 and 5/9 golden dataset ion-GCF pairs observed above a 1% FDR at 0.30 and 0.50 glocal similarity score cutoffs, respectively.

Based on the BGCs and molecules observed in the above data, gene cluster networks and molecular networks were generated. Exploration of these networks highlighted high diversity in both gene clusters and molecules, exemplified by the identification of 152 different BGCs (at <0.95 overall similarity) related to known detoxin/rimosamide gene clusters (Fig. 3b), and 110 different molecules related to detoxins and rimosamides (Fig. 3c).

**Fig. 3.**
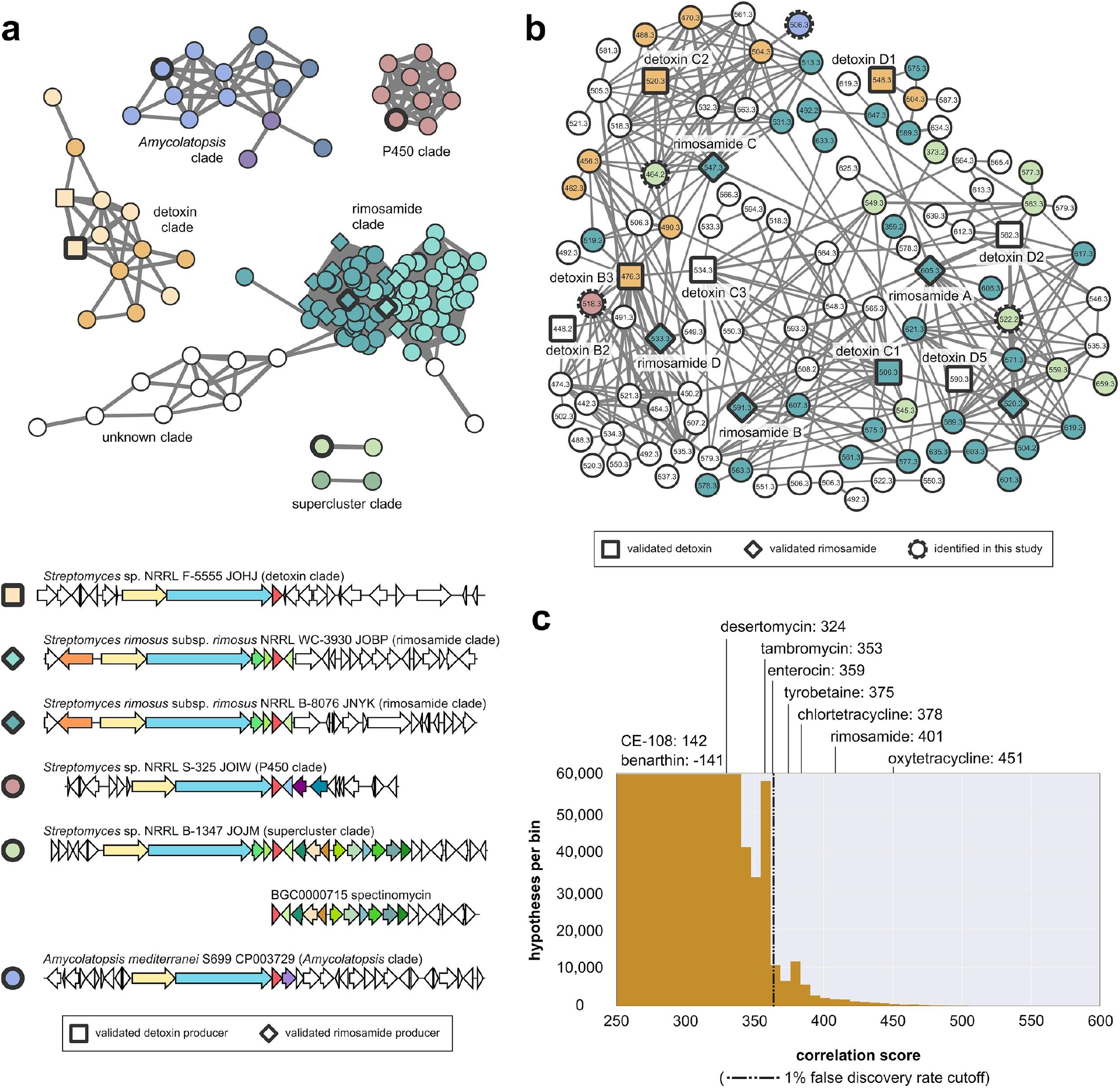
**a,** Detail of a BiG-SCAPE NRPS BGC network (filtered for the presence of the TauD domain) containing BGCs related to the known gene clusters for detoxin/rimosamide biosynthesis. Color shades reflect BiG-SCAPE family classification related to the colored clades in Fig. 5, i.e. families with a rimosamide producer (turquoise), with a detoxins producer (orange), and two additional abundant families named the *Amycolatopsis*/P450 clade (violet), P450/enoyl clade (green) and the spectinomycin ‘supercluster’ clade (light green). Representative clusters for each clade are shown at bottom with the spectinomycin gene cluster from MIBiG displayed for reference. Gene colors reflect those in the CORASON tree of Fig. 5. Importantly, not all nodes in this families contain clusters from the high-resolution clades of the CORASON tree due to the fragmented nature of the assemblies and the presence of relatively close Core Biosynthetic Genes detected by antiSMASH (see Fig. S10). **b,** Molecular network of tandem MS data from a 363 strain actinomycete library is colored by BiG-SCAPE clade and includes 103 putatively novel detoxins and rimosamides. Known detoxin and rimosamide nodes are squares and diamonds, respectively, with solid bold outlines while novel detoxins detected from *Streptomyces spectabilis* NRRL 2792 (*m/z* 464.240, *m/z* 522.247 in light green) and *Amycolatopsis jejuensis* NRRL B-24427 (two isomers with *m/z* 506.287 in violet) are indicated by bold dotted outlines. **c,** Histogram of all ion-GCF correlation scores resulting from a metabologenomics round run with 0.30 glocal similarity score cutoffs. Known ion-GCF pair correlation scores are overlaid, with tyrobetaine, chlortetracycline, rimosamide, and oxytetracycline ion-GCF pairs identified above a 1% FDR threshold.

### Multi-locus gene cluster phylogenies resolve evolutionary relationships between related BGCs

Genetic diversity of BGCs within GCFs is directly related to structural differences between their molecular products, and even small chemical variations can lead to different biological activities^37^. Hence, mapping the evolutionary relationships between BGCs within and across GCFs is crucial for the discovery process. To this end, we introduce the *CORe Analysis of Syntenic Orthologues to prioritize Natural products biosynthetic gene clusters* (CORASON) software, written in Perl and available as open source from https://github.com/nselem/corason. Given a query gene inside a BGC of interest, the CORASON pipeline identifies other genomic loci that contain homologues of this gene and calculates a multi-locus phylogeny of all loci based on their conserved core (Fig. 4).

**Fig. 4.**
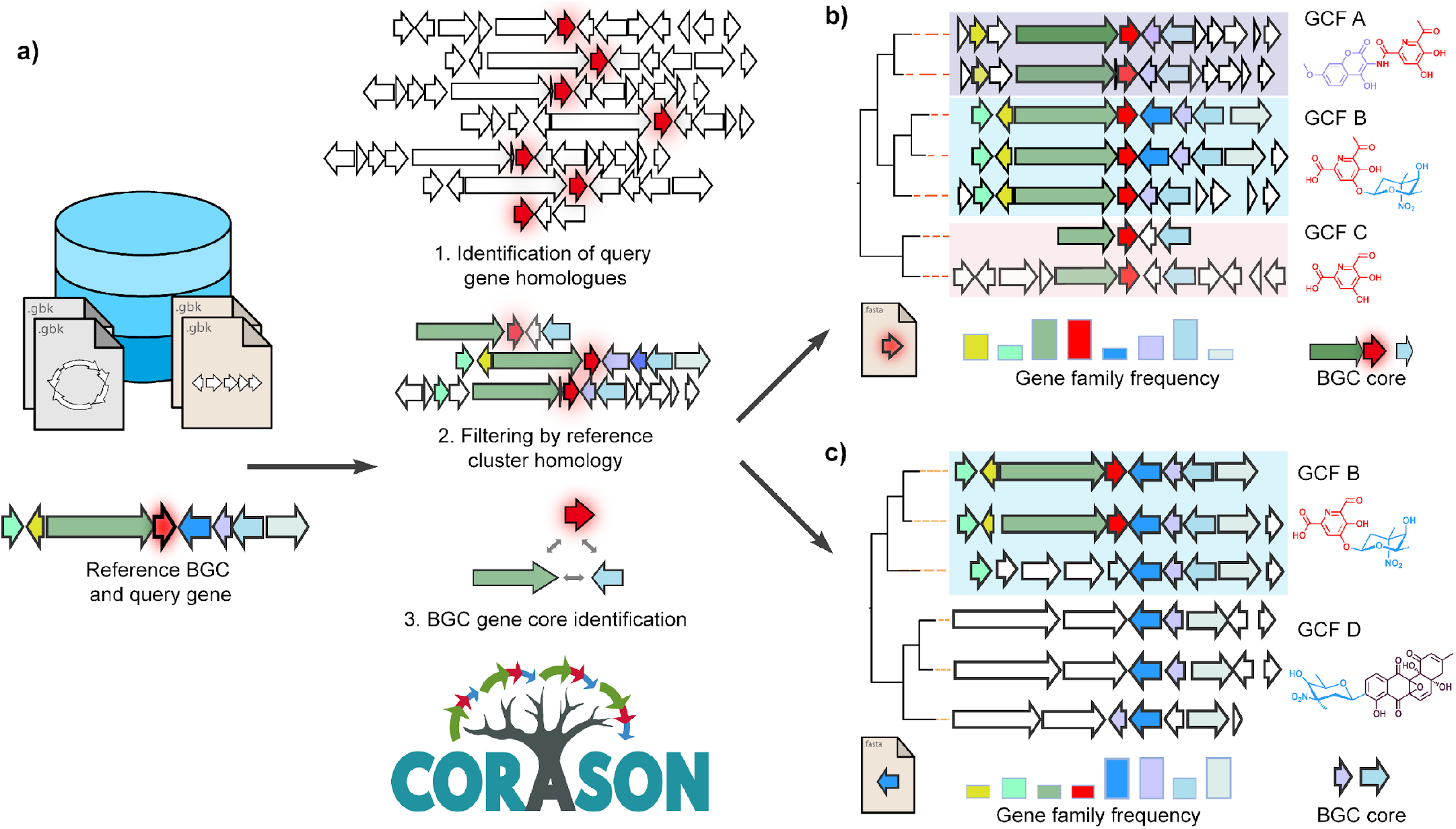
CORASON Workflow. **a,** Given a query gene in a reference cluster and a custom genomic annotated database, CORASON i) searches for query gene homologues, ii) creates the Cluster Variation Database (CVD) by filtering out all genomic vicinities not related to the reference BGC, including keeping fragmented clusters and iii) calculates the CVD gene core by multi bidirectional best hits. **b,** Then, CORASON infers a phylogenetic tree by curation and concatenation of the CVD gene core, and displays the CVD following the tree and calculates the frequency of occurrence for each gene family from the reference BGC. **c,** With the same reference BGC, if a new query gene is selected from accessory enzymes instead of the current CVD core, CORASON will visualize a new phylogeny with families containing the same molecular modifications, expanding the chemical universe within an evolutionary framework.

CORASON is available for users as a downloadable and easy-to-install software that allows tracing the evolutionary history of biosynthetic genes using customizable databases. Any subset of genes from a BGC may be selected to identify other genomic contexts across a set of prokaryotic genomes in which homologues of these genes are found. The selected query genes are then visualized on a multi-locus approximate-maximum-likelihood phylogenetic tree^46^ that allows the user to identify all functional genomic contexts in which the corresponding gene families are found. In this way, the evolutionary relationships of each gene within a BGC across large numbers of genomes can be comprehensively analyzed. A version of the CORASON algorithm, called ‘family-mode’, was also integrated with BiG-SCAPE, allowing to generate a multi-locus phylogeny of all BGCs within each GCF using the sequences of their common domain core.

### An integrated workflow and interactive visualization allow rapid exploration of biosynthetic diversity

BiG-SCAPE and CORASON connect seamlessly with antiSMASH and MIBiG, as GenBank outputs of antiSMASH can be used directly as inputs for the workflow, and MIBiG reference data can be included in the analysis automatically. Although calculations on hundreds or thousands of genomes are too compute-intensive to provide them on a free public web server, we still wanted to make the results available in an interactive user-friendly HTML visualization that enables efficient exploration of biosynthetic diversity across large datasets for non-programmers. Hence, we constructed a powerful JavaScript-based visualization that provides an interactive output for every BiG-SCAPE run, which can be viewed offline on any web browser. In a single view, the visualization displays BGC nodes colored by GCF in interactive sequence similarity networks, side-by-side with arrow visualizations of the gene clusters, which contain gene annotation and Pfam domain details that appear on mouse-over. Networks can be searched by compound names of MIBiG reference clusters, Pfam domains of interest, or species names, with resulting match nodes instantly highlighted within the network. Each GCF is given its own view panel, which shows the CORASON-based multi-locus phylogeny of the underlying BGCs and includes links to related families within the same GCF. Finally, an overview page is provided that displays statistics on the BGCs identified, as well as a GCF absence/presence heatmap of the most frequently occurring gene clusters within the input dataset.

To illustrate BiG-SCAPE/CORASON usage, we provide an example output of a run with antiSMASH-predicted BGCs from 103 complete *Streptomyces* genomes, including as outgroups the genomes of *Catenulispora acidiphila* and *Salinispora arenicola*: http://bioinformatics.nl/~xnava009/streptomyces_out/. To connect the absence/presence map of GCFs across these genomes to species phylogeny, a high-resolution multilocus whole-genome phylogeny (Fig.S11) was inferred from the *Streptomyces* conserved-core (Online Data: StreptomycesCore.), and the GCF absence/presence patterns were plotted onto the structure of the tree (Fig. S12). As has been observed before in other genera like *Salinispora*^29^, this shows high conservation of select GCFs across larger numbers of genomes, combined with large numbers of rare GCFs that are specific to one or a few genomes.

### Identification of novel detoxin/rimosamide analogues using BiG-SCAPE and CORASON

To showcase the power of our workflow for the analysis of large BGC collections and high-resolution mapping of GCF biosynthetic diversity, we focused on the detoxin and rimosamide GCFs^38^. Our comprehensive analysis of the selected actinobacterial genomes revealed these BGCs to be taxonomically widespread and architecturally diverse (Fig. 5). The conserved gene core of detoxin and rimosamide BGCs is composed of three genes: one NRPS, one NRPS/PKS hybrid, and a homologue of *tauD*, presumably recruited from the *tauABCD* operon in *Escherichia coli*^39^. as suggested by EvoMining analysis.

**Fig. 5.**
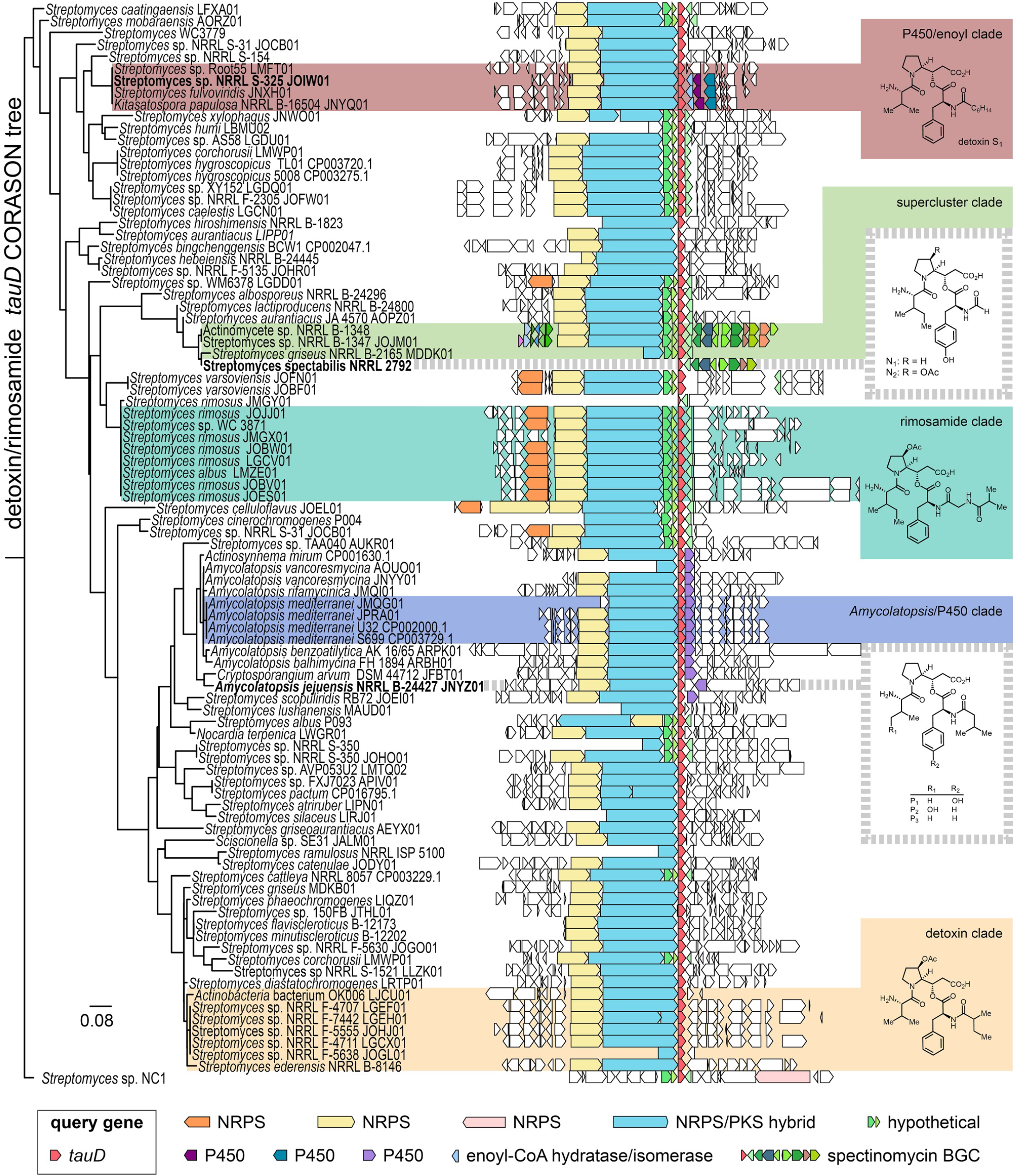
CORASON phylogeny of detoxin/rimosamide-related BGCs. CORASON phylogenetic reconstruction with *tauD* as query gene and the *Streptomyces* sp. NRRL B-1347 BGC as query cluster. Some highly similar BGCs and those that were the most remote from the BiG-SCAPE-defined clades were removed from the original tree for readability. The representative structures for each clade illustrate the correspondence between molecular and genomic variations. The BGC tree is rooted with a TauD encoded in an unrelated (non-NRPS) BGC from *Streptomyces* sp. NC1. The genes encoding the NRPS, the NRPS/PKS hybrid, and the TauD homologue are highly conserved among detoxin and rimosamide BGCs. Highlighted sections on the tree indicate BiG-SCAPE-defined clades. Bolded strain/BGC names are those investigated in this study with dotted lines indicating those which were just outside the BiG-SCAPE-defined clades. Clusters in the ‘P450/enoyl clade’ contain a pair of P450 genes and an enoyl-CoA hydratase/isomerase which are putatively involved in biosynthesis of alkylated detoxins, the ‘supercluster clade’ is comprised of detoxin clusters and is immediately adjacent a spectinomycin BGC, and the ‘*Amycolatopsis*/P450 clade’ possess a unique P450 enzyme that corresponds to the first examples of hydroxylation in the detoxin/rimosamide class. Strain names are followed by their Genbank accession number when available. Genes not found in the reference cluster are colored based on BLAST analysis.

The rimosamide BGC differs from those of the detoxins by having an additional NRPS, which encodes for further elaboration of the common detoxin/rimosamide core scaffold with isobutyrate and glycine^38^. Aside from the more common NRPS and PKS genes, the fact that the *tauD* gene was present across all members of this family caught our attention. The product of the *tauD* gene belongs to the Fe(II)/α-ketoglutarate-dependent hydroxylase enzyme superfamily and is named for the commonly encoded α-ketoglutarate-dependent taurine dioxygenase involved in assimilation of sulphite by oxygenolytic release from the amino acid taurine^40^. Interestingly, this family also includes enzymes across fungi, bacteria and plants that catalyze hydroxylations, desaturations, ring expansions, and ring formations, among other transformations. To date, the role of *tauD* in detoxin and rimosamide biosynthesis is unknown, but it has been suggested to be responsible for the proline oxidation observed in some analogs^38^.

An EvoMining^8^ analysis of the TauD dioxygenase protein family showed specific specialized metabolism-related expansions of paralogues across genera such as *Streptomyces, Rhodococcus*, *Frankia* and *Amycolatopsis* (Fig S13). Within this expansion, one clade was shown to contain fifteen *tauD* homologues that are part of experimentally characterized BGCs from MIBiG, including the detoxin and rimosamides BGCs (Table S6).

To identify novel detoxin and rimosamide-related natural product BGCs, we selected all BGCs containing a *tauD* homologue. The resulting 1175 gene clusters were then subjected to a combined BiG-SCAPE/CORASON analysis. This revealed that the detoxin and rimosamide GCFs are part of a larger gene cluster clan related to peptide biosynthesis that also comprises unexplored families across the phylum Actinobacteria (Fig 5). Metabolomic data was available for 40 of the 152 strains identified to encode these BGCs. Tandem mass spectrometry molecular networking analysis of these strains indicated the presence of three known detoxins, four known rimosamides, and 103 putatively novel detoxin and rimosamide analogues (Fig. 3b), confirming the vast natural product chemical diversity suggested in the BiG-SCAPE/CORASON data.

Based on inspection of genetic features unique to detoxin/rimosamide BGC phylogenetic clades identified by CORASON (Fig. 5, in red), there were three that captured our interest. The first was named the ‘P450/enoyl clade’ and contained two P450 genes and an enoyl-CoA hydratase/isomerase within its BGCs. Tandem mass spectrometry analysis of extracts from *Streptomyces* sp. NRRL S-325, which contains a BGC within this clade, led to the discovery of detoxin S_1_ (**1**; Figs. 5, S15-16) with a heptanamide side chain, a unique feature among the detoxins and rimosamides whose installation likely depends on the enoyl-CoA hydratase/isomerase.

The second clade of interest, termed the ‘supercluster clade’ (Fig. 5, in light green), comprised BGCs that encode detoxins in a ‘supercluster’ that included the known spectinomycin BGC. Two detoxin-like BGCs within this clade in the genomic dataset, identified in the genomes of *Streptomyces* sp. NRRL B-1347 and *Actinomycete* sp. NRRL B-1348, were syntenic to a spectinomycin BGC (virtually identical to BGC0000715 in MIBiG) in a supercluster configuration (Fig. 5). Interestingly, the MIBiG spectinomycin cluster was included on the detoxin/rimosamide CORASON tree just beside the BiG-SCAPE-defined supercluster clade due to the presence of a *tauD* gene at its periphery that is homologous with those from the detoxins. The *tauD* gene is not known to be involved in spectinomycin biosynthesis, so we hypothesized that the strain from which the MIBiG entry was sourced, *Streptomyces spectabilis* NRRL 2792, must have a detoxin BGC beside the spectinomycin cluster and would also produce a detoxin-like product. With this, we acquired the producing strain to determine if CORASON analysis was powerful enough to predict metabolite production solely based on the presence of a query gene but in the absence of full genomic data or a complete BGC. Tandem mass spectrometry analysis of a *S. spectabilis* NRRL 2792 extract revealed production of six detoxin-like natural products, including detoxin N_1_ (**2**; Figs. 5, S17) and its acetoxylated analog, detoxin N_2_ (**3**; Figs. 5, S19). Structural features unique to this novel detoxin subclass included the incorporation and formylation of tyrosine resulting in a formamido group. LC-MS analysis of cultures supplemented with stable isotope–labeled amino acids corroborated these structural predictions based on analysis of the BGC and tandem MS data (Figs. S18, S20).

The third detoxin clade that we targeted occurred almost entirely within the genus *Amycolatopsis* and was named the ‘*Amycolatopsis*/P450 clade’ (Fig. 5, in blue). This clade drew our interest as its BGCs contained a cytochrome P450 gene unique among the detoxins and rimosamides that was predicted to be responsible for novel hydroxylation patterns. Though we did not have strains in our library with BGCs in the BiG-SCAPE-defined clade, CORASON analysis allowed the selection of a strain with a very similar BGC also containing the desired P450 gene (Fig. 5). Tandem MS analysis of this strain, *Amycolatopsis jejuensis* NRRL B-24427, revealed detoxin isomers P_1_ (**4**; Figs. 5, S21) containing a tyrosine, P_2_ (**5**; Figs. 5, S26) featuring a hydroxylated valine, as well as detoxin P_3_, a closely related analog free of hydroxylation (**6**; Figs. 5, S31). As before, validation of amino acid assignments observed in MS^2^ fragmentation data was achieved through several metabolic feeding experiments using stable isotope-labeled amino acids (Figs. S22–S25, S27–S30, and S32). Detailed structural analysis of **1–6**, including detailed interpretation of tandem MS spectra and results from feeding studies using stable isotope-labeled amino acids, can be found in the Supporting Information.

Our results show how BiG-SCAPE can effectively identify sets of related BGCs across large numbers of genome sequences. Moreover, using CORASON to systematically map BGC evolutionary diversity and assemble gene cluster phylogenies proved to be a powerful approach for the discovery of novel clades of BGCs that are responsible for the biosynthesis of uncharted natural product chemistry. When focused toward the specific detoxin/rimosamide discovery effort in “query mode,” CORASON guided the discovery of six new detoxins by way of *tauD* phylogeny. Thus, CORASON enables rapid mapping and visualization of BGC relationships for more effective genome mining from large genomic libraries. Additionally, variation in BGC domain architecture corresponded to variation in chemical structure – presence of an enoyl-CoA hydratase/isomerase corresponded to the fatty acid amide detoxin S_1_ and presence of a unique P450 gene corresponded to evidence of hydroxylations in detoxins P_1_–P_3_.

## Conclusions and future perspectives

The comprehensive computational workflow introduced here enables effective exploration of biosynthetic diversity across large strain collections, pan-genomes of entire bacterial or fungal genera, and metagenomic datasets with thousands of metagenome-assembled genomes. The combined BiG-SCAPE/CORASON platform overcomes key computational bottlenecks inherent in previous approaches, as it facilitates building GCFs with both partial and complete BGCs, accounts for class-specific differences between BGCs, incorporates sequence identify information within limited compute time, charts out evolutionary relationships between and within GCFs, and provides an interactive user interface to explore the outputs. Hence, we anticipate that it will soon facilitate metabologenomic correlation studies to systematically assign many gene clusters to many molecules at unprecedented scales.

Additionally, the ability to perform phylogenetic analyses of large sets of complete BGCs, as well as their individual genetic components, a long-standing challenge that has remained unsolved since first posed by Fischbach, Walsh and Clardy in 2008^41^, will constitute a key technology to facilitate fundamental studies on the evolutionary origins of natural product chemical innovations. For example, it provides a stepping-stone to perform detailed analyses of how gene cluster architectures have evolved (and are evolving) from their constituent independent enzymes and sub-clusters. A logical next step will be the unified classification of the millions of BGCs within publicly available genome sequences, and a Pfam-like database for the assignment of biosynthetic gene cluster families to known and unknown areas of natural product chemical diversity.

## Acknowledgements

We express our gratitude to the Agricultural Research Service of the United States Department of Agriculture for providing bacterial strains; Hyoung Sook Ann, Zachary Crispino, Yewool Kim, Natalia Ciszek, and Kyle Espejo for generating bacterial culture extracts; Anthony Goering, Ryan McClure, Matthew Robey, and Galen Miley for assistance with and contributions to metabolomic data collection methods and acquisition. Some analyses were carried out using CONABIO’s computing cluster, with funds from SEMARNAT.

The research reported in this publication was supported by the Netherlands Organization for Scientific Research (NWO) VENI Grant [863.15.002 to MHM], the Graduate School for Experimental Plant Sciences (EPS grant to MHM); National Institutes of Health Genome to Natural Products Network supplementary award [U01GM110706 to MHM], CONACyT [grants CBS2017_285746 and 2017_051TAMU to FBG]; postdoctoral scholarship 263661 to JCNM,; PhD scholarship 204482 to NSM, who was also supported by Innovation Secretary of Guanajuato), the National Cancer Institute (NCI) of the National Institutes of Health (NIH) under Award Number F32CA221327 (MWM), the National Institute of General Medical Sciences (NIGMS) under Award Number F32GM120999 (EIP), the National Center for Complementary and Integrative Health (NCCIH) of the NIH under Award Number R01AT009143 (RJT, NLK).

## Online Data

Online data is publicly available at DOI: 10.5281/zenodo.1237237

## Online Methods

### BGC network preparation

A set of 2,831 Actinobacterial genomes was downloaded from NCBI by querying for "Whole genome shotgun sequencing project" or "Complete genome" in combination with the taxonomic identifier for actinobacteria. The Propionibacteriales, Micrococcales, Corynebacteriales and Bifidobacteriales orders were excluded, as they contain large numbers of genomes without relevant natural product-producing capacity, with the exception of the Nocardiaceae family from the Corynebacteriales. To these set, 249 additional unpublished draft genomes from the Metcalf lab were added. All files were processed with antiSMASH v4^42^ (no ClusterFinder or other special parameters enabled). Antismash-processed genome sequences are available as Online Data (antiSMASH_results_Metcalf_B, antiSMASH_results_Metcalf_J and antiSMASH_results_NCBI).

To the resulting 73,260 predicted Biosynthetic Gene Clusters (BGCs), 1,393 more were added from the Minimum Information about a Biosynthetic Gene Cluster database (MIBiG^20^, release 1.3, August 2016, antiSMASH-analyzed versions from each individual entry) as reference data. These final BGC set was then analyzed with BiG-SCAPE using version 31 of the pfam database. The `hybrids` mode, which allows BGCs with mixed annotations be analyzed in their individual Class sets (e.g. a BGC annotated as *lantipeptide-t1pks* will be analyzed as both a RiPP and a PKSI) was enabled. Two results sets were created, one with the default "global" mode enabled, and the other with "glocal" mode enabled (See Fig. 2)

The CORASON and EvoMining results used the same unpublished draft genomes but a reduced set of 1,668 Actinobacterial genomes from an older query on the NCBI website. (see Text S5).

### Alignment Method Comparison

To compare alignment methods for domain sequences, the normal version of BiG-SCAPE was used (snapshot https://git.wur.nl/medema-group/BiG-SCAPE/commit/992b0c6a3c0a35ac46d56227018d5d17d5b7e789) against a custom-prepared version of the same snapshot using Muscle 3.8.1551^43^ [Online Data: AlignmentMethodComparison]. Both versions used default parameters and the --no-classify flag (so only sequence extraction, domain prediction and domain alignment were performed). The hmmalign version took 923s whereas the Muscle version took 13392s (3h:43m:12s) on a desktop PC with Intel Core i7-7700 CPU and 16 GB of RAM.

### Clustering algorithm optimization

The election of the clustering algorithm was based on an initial analysis of the BGCs from the MIBiG database using BiG-SCAPE (--hybrid mode disabled). In this analysis, the network went through a targeted attack in order to first identify the most suitable cutoff for clustering algorithm evaluation (Fig. S2). Networks in terms of number of vertices/edges lost for each cutoff value, as well as the size of the connected components that emerged were calculated during the attack. c=0.75 was chosen as it maximized the number of nodes while minimizing the impact to the network structural integrity. Using this cutoff value, entropy was calculated for several clustering algorithms (Text S4, Figs. S3 and S4) using the Curated Compound data [Table S1]. The Affinity Propagation clustering method showed the most sensible results, producing clusters with low entropy and average size. See further details in Text S3.

### BiG-SCAPE algorithm implementation summary

#### Input data set

The input of BiG-SCAPE are text files in GenBank format (.gbk extension, ideally processed with antiSMASH) as well as the Pfam database (already processed with hmmpress).

#### Algorithm overview

After selecting and filtering (e.g. for certain size, in base pairs) the input GenBank files, protein sequences are extracted. All the sequences from each file are searched for conserved domains using a user-supplied external Pfam database. Overlapping domains are filtered based on the score calculated by hmmer. The sequences of every predicted domain type are aligned using each corresponding model by hmmalign. A distance matrix is created by calculating the distance between every pair of BGC in the data set.

#### Distance calculation

Pairwise distance calculation is divided between three values that measure a) percentage of shared domain types (Jaccard Index), b) Similarity of aligned domain sequences (Domain Sequence Similarity index; domains from the same type are first matched for best similarity using the Munkres algorithm, as implemented in Scikit-Learn library^44^) and c) Similarity of domain pair-types (Adjacency Index). For specific details of each index, see Text S1.

There are two ways of selecting the domains predicted within each BGC for the calculation of distance. In the global mode, all domains are considered. For cases where difference in size is large (due to e.g. one BGC being placed at the edge of a contig or when comparing curated BGCs with shorter gene borders), we implemented the so-called glocal mode, where a selection of domains is used in the distance calculation. In this mode, genes in each BGC are represented as a concatenated string of Pfam domains, and each BGC in the pair is represented as a list of those domain concatenations (strandedness is not taken into account).

BiG-SCAPE then uses the SequenceMatcher method from Python’s difflib library to to find the longest match (internally called the LCS or "Longest Common Subcluster"). The second BGC is tried in the reverse orientation and the orientation with the largest LCS is kept.

To proceed to the next step, the LCS must be either three genes long, or contain at least one gene marked by antiSMASH as "Core Biosynthetic".

In the extension stage the selection of domains is extended for the BGC with least number of genes up(down)stream. The remaining BGC domain selection (per side) will be tried for expansion according to the following scoring algorithm in the Alignment Stage: for every gene in the reference BGC, a gene with the same domain organization is searched for in the remaining BGC. If such a gene is found, the score will be added a bonus (match=5) plus a penalty proportional to the distance from the current position (number of genes * gap penalty where gap=-2) and the current position will be moved to the position of the matching gene. If a gene with the same domain organization is not found, the score will be decreased with a penalty (mismatch=-3).

#### GCF clustering

Once the distance matrix is calculated for each BiG-SCAPE class (see Table S5), Gene Cluster Family (GCF) assignment is performed for every cutoff distance selected by the user (the interactive visualization of BiG-SCAPE will show the one with the largest number) with 0.3 being the default. For every cutoff, BiG-SCAPE creates a network using all distances lower or equal than the current cutoff. The Affinity Propagation clustering algorithm^34^ is applied to each subnetwork of connected components that emerge from this procedure. The similarity matrix for Affinity Propagation includes all distances between members of the subnetwork (i.e. it includes those with distance greater than the current cutoff).

Gene Cluster Clan (GCC) setting (enabled by default) will perform a second layer of clustering on the GCFs. For this, Affinity Propagation will be applied again but network nodes are represented by the GCFs, defined at the cutoff level specified in the first value of the --clan_cutoff parameter (Default: 0.3). Clustering will be applied to the network of all GCFs connected by a distance lower or equal than the GCC cutoff (second value of the --clan_cutoff parameter; larger distances are discarded. Default: 0.7). Inter-GCF distance is calculated as an average distance between the BGCs within both families. Affinity propagation parameters: damping=0.9, max_iter=1000, convergence_iter=200.

#### Output

BiG-SCAPE produces high-quality SVG figures for every BGC as well as text files from each of its algorithm (hmmer domtable results, filtered domain results, aligned domain sequences, clustering results and the distance network). As part of the output, BiG-SCAPE also offers an interactive visualization where the user can see an overview of the distance network generated by the highest cutoff selected. BGCs connected and clustered into GCFs have a page on their own for closer inspection.

#### CORASON-like GCF tree

As part of BiG-SCAPE’s visual output, a CORASON-like tree is generated for every GCF page. This tree is created using the sequences of the Core Domains in the GCF. These are defined as the domain type(s) that a) appear with the highest frequency in the GCF and b) are detected in the central (or “exemplar”) cluster, defined by the Affinity Propagation cluster. All copies of the Core Domains in the exemplar are concatenated, as well as those from the best matching domains of the rest of the BGCs in the GCF (aligned domain sequences are used). The tree is constructed using FastTree (default options). Alignment is attempted using the position of the Longest Common Information from the distance calculation step (between the exemplar BGCs and each of the other clusters)

#### Availability

BiG-SCAPE is written in Python and currently compatible with Python 2 and Python 3. It is freely available at https://git.wageningenur.nl/medema-group/BiG-SCAPE. More extensive details of the algorithm are available at the repository’s wiki: https://git.wageningenur.nl/medema-group/BiG-SCAPE/wikis/home

### Weight optimization methods

Tuning of weights for each BiG-SCAPE class was calculated by a brute-force approach, by choosing the weight combination that maximized the correlation between BGC and Compound distances for every pair of BGCs in the same class in a manually curated Compound Group table (Table S1). The data set comprised all BGCs from the MIBiG database (v1.3) that had linked compound SMILES and had at least 2 predicted domains to filter out minimal gene cluster entries. BGC distances were calculated by moving in steps of 0.01 between the Jaccard, Domain Sequence Similarity, the original Goodman-Kruskal^45^, and Adjacency indices, such that JI+DSS+GK+AI=1. The anchorboost parameter of DSS was allowed to change in the range [1,4] with steps of 0.5. For the DSS index, only the original 4 anchor domains were considered (Condensation Domain, PF00668; Beta-ketoacyl synthase N-terminal, PF00109; Beta-ketoacyl synthase C-terminal domain, PF02801 and the Terpene synthase N-terminal, PF01397). Compound distances were calculated only once, between all BGCs in the MIBiG 1.3 that had an annotated SMILES string representing the molecule. Their pairwise distance was calculated by using RDKit (Tanimoto coefficient based on Morgan fingerprinting, radius=4). The nine original curated Compound classes was used to tune the weights of 7 BiG-SCAPE classes (the Terpene BiG-SCAPE class was originally included in the Others Compound class due to a low number of points and was assigned default values of Jw = 0.2, DDSw = 0.75, AIw = 0.05).

Results indicated clear tendencies to favor different indices in each case, and corroborated that the proposed Adjacency Index was more significant than the original Goodman-Kruskal synteny metric (additional details in Text S2 and Fig. S1).

### Methods for CORASON algorithm

CORASON inputs are a custom genomic database, a reference cluster and a query gene located within the reference cluster. The genomic database is a collection of either genomes or BGCs in GenBank format. CORASON will identify the conserved core of the reference BGC within the genomic database. To calculate the conserved core, a generalization of best bidirectional hits (BBH) comparison was used. The BBH relationship is a property among two sets of genes, that can genomes, metagenomes or as in this case BGC. This relationship was generalized in a stricter algorithm that consideres instead of two BGC, every BGC in a collection. As a result, in this particular case, families of the conserved core are composed only by genes that are BBH of the corresponding orthologue for every BGC in the genomic database. This strict criterium is needed to remove paralogues and to conserve only real orthologues. Multidirectional best hits are those genes in a list that are best bidirectional hits all versus all, in contrast with best bidirectional hits were only pairs of genes are compared. The BGC gene core facilitates reconstructing the BGC evolutionary history in a multilocus tree. The query gene assures that at least one element will be present in the conserved core. The query gene will also be used to visually align the BGCs in the graphical output.

### Identification of reference BGC variations on the genomic DB

CORASON uses BlastP, with an E-value cutoff of 0.001 to find all query enzyme homologues within the genomic database. The genomic contexts of the query enzyme homologues are expanded 10 genes at each side and stored in a temporary database. Next, sequences from the reference BGC located within less than n genes (default: n=10) from the query enzyme are blasted against the temporary database using the same E-value cutoff. Genomic context size, E-value and bit score cutoffs are user-adjustable parameters. Finally, all genomic contexts with at least two homologues, including the query enzyme and at least one additional homologue from the reference cluster, are kept as the Cluster Variations Database (CVD) for further analysis.

### Gene core determination

To reconstruct the phylogeny of the BGC variations, the conserved core is calculated. The core is strongly dependent on the taxonomic diversity of the organisms considered and also on the genome quality. For instance, if the BGCs are not closely related, the core can be reduced to only the query gene. A set of homologous genes are considered part of the conserved-core if and only if they are shared among the cluster variations internal database (all BGCs) and are multidirectional best hits i.e. if they are best n-directional hits in an all vs all manner.

Within a set of *N* BGCs variations, if the set *H* of homologous genes is defined as follows: 
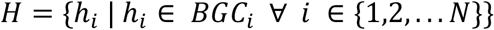
then, *H* belongs to the conserved core if and only if 
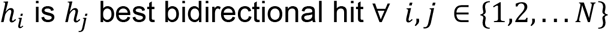

### Phylogenetic reconstruction and gene cluster alignment

For each BGC, its conserved core sequences are concatenated and then aligned using Muscle version 3.8.31 ^43^. The alignments are curated using Gblocks ^48^ with a minimum block length of 5 positions, a maximum of 10 contiguous nonconserved positions and considering only positions with a gap in less than 50% of the sequences in the final alignment. If the curation turns out to be empty, then the noncurated alignment will be used for the tree. If the alignment itself is empty, it is recommended to the reduce the score cutoff or the scope of the taxonomic diversity on the genomic database. Without the alignment, BGCs will be drawn but not sorted. Approximately-maximum-likelihood phylogenetic trees are inferred using FastTree ^46^ version 2.1.10 from the curated amino acid alignment.

### BGC prioritization graphical output

CORASON produces a Scalable Vector Graphics (SVG) file containing the BGC variations sorted as stated by the phylogenetic reconstruction and aligned according to the query enzyme. The newick tree is converted to SVG applying Newick Utilities version 1.6 ^47^ and each BGC is drawn with the Perl module SVG. As an additional feature to facilitate even more visual differentiation of BGC families within BGC clans, genes on each cluster are visually represented with a color gradient according to the sequence similarity to their homologous gene on the reference cluster. Other CORASON outputs include the Newick tree, the genbank files of the BGC variations and the conserved core report.

CORASON was developed on Perl:5.20 and is available as free software on GitHub (https://github.com/nselem/corason) and as a downloadable image on dockerhub (https://hub.docker.com/r/nselem/corason/). A CORASON tutorial is available on-line at https://github.com/nselem/corason/wiki.

### Methods of BGCs families on *Streptomyces* dispensable genome

Sequences from 103 complete *Streptomyces* genomes were retrieved from NCBI by querying for "*Streptomyces*" and "complete genome" not “segment”. The conserved core was extracted and curated with the CORASON algorithm. The tree was constructed using FastTree with default values over a matrix of 114,051 amino acids in size, from 446 conserved gene families(Online Data: StreptomycesCore).

### Phylogenomic analysis

For the TauD expansions tree (Fig S13), a *tauD* sequence from *Escherichia coli K12* as query to conduct a blast search against the reduced genomic database of 1917 Actinobacteria genomes (e-value .001), followed by an EvoMining analysis and a search for recruitments on MIBiG database (e-value 0.001). Recovered *tauD* orthologues were aligned with Muscle 3.8.31 ^43^ and alignments were curated using Gblocks ^48^ in the same manner as described above. An unrooted Approximately-maximum-likelihood was built using FastTree^46^, an algorithm specialized on very large protein families. Tree was colored using Newick Utilities^47^ according to BiG-SCAPE families.

The CORASON tree has as query enzyme TauD from the reference cluster of the organism *Streptomyces NRRL* B-1347 (Accession JOJM01). CORASON trees are unrooted, but this tree was posteriorly rooted with the BGC from the genome *Streptomyces sp NC1*, because this BGC is different from all other clusters in the dimeric peptide clan, as it does not share the core but the accessory enzymes with other BGC clan members.

